# Transitive inference in humans and rhesus macaques after massed training of the last two list items

**DOI:** 10.1101/055335

**Authors:** Greg Jensen, Yelda Alkan, Fabian Muñoz, Vincent P. Ferrera, Herbert S. Terrace

## Abstract

Transitive inference (TI) is a classic learning paradigm for which the relative contributions of experienced rewards and representation-driven inference have been vigorously debated, particularly with regard to the notion that animals are capable of logic and reasoning. Rhesus macaque subjects and human participants performed a TI task in which, prior to learning a seven-item list ABCDEFG, a block of trials presented exclusively the pair FG. Contrary to the expectation of associative models, the high prior rate of reward for F did not disrupt learning of the entire list. Monkeys (who each completed many sessions) learned to anticipate that novel stimuli should be preferred over F. We interpret this as evidence of a general task representation of TI that generalizes beyond learning about specific stimuli. Humans (who were task-naïve) showed a transitory bias to F when it was paired with novel stimuli, but very rapidly unlearned that bias. Performance with respect to the remaining stimuli was consistent with past reports of TI in both species. These results are difficult to reconcile with any account that seeks to assign the strength of association between individual stimuli and rewards. Instead, they support both sophisticated cognitive processes in both species, albeit with some species differences.

## 1 Introduction

Transitive inference (TI) is a fundamental process in propositional logic, and has been studied by psychologists for over a century (Burt, 1911). If A > B, and B > C, and all three items belong to an ordered list, then the transitive property of the “>” operator permits the conclusion that A > C. Thus, TI provides a formal definition for what it means to “know” that a set of items are ordered.

The first animal study of TI was reported by McGonigle & Chalmers (1977). They presented squirrel monkeys (*Saimiri sciureus*) with adjacent pairs of stimuli from a five-item list (i.e. for a list of stimuli ABCDE, subjects were initially trained on AB, BC, CD, and DE). The “correct” choice was whichever item came earlier in the list. Once subjects reliably selected correct items, they were tested on all ten possible pairs. Despite having never seen the nonadjacent pairings previously, subjects not only selected the correct items with high accuracy, but did so at comparable rates to 4-year old human children.

Since this initial study, accurate performance on TI tasks has steadily been reported in a growing range of species, and the capacity for TI in animals appears to be ubiquitous among vertebrates (Vasconcelos, 2008). Some form of TI has been reported in at least 20 species (Jensen, In Press), suggesting that TI taps into cognitive faculties that are well preserved across a wide evolutionary range of species.

Two major behavioral features have regularly been reported in the TI literature. One is the terminal item effect (Wynne, 1997). The first item in a TI list is always rewarded, while the last is never rewarded, so these items are less ambiguous than the non-terminal items. This appears to yield a boost to their accuracy. The other is the symbolic distance effect (D'Amato & Colombo, 1990). Adjacent pairs typically evoke the lowest response accuracy, whereas items two positions apart (e.g. BD) have higher accuracy, and so on such that the pair comprising the two terminal items has the highest accuracy. In both cases, higher accuracy tends to be correlated with lower reaction times, so terminal item effects and symbolic distance effects are sometimes also reported with respect to reaction time (McGonigle & Chalmers, 1992).

Despite this rich literature of empirical results, the field remains divided on how TI itself is performed. The cognitive approach to explaining TI in animals rests on the premise that subjects form a representation of the list (Zentall, 2001; Jensen et al., 2015). Several lines of evidence are consistent with the cognitive approach. For example, subjects are able to transfer prior serial learning between TI and a serial task with qualitatively different task demands (Jensen et al., 2013). In addition, the serial position of each stimulus can be dissociated from its reward magnitude without disrupting subsequent TI (Gazes et al., 2012).

Many studies of TI in animals reject the cognitive interpretation in favor of models of associative learning (Couvillon & Bitterman, 1992; Allen, 2006). Under the logic of this approach, subjects are able to infer the order of stimuli by determining how strongly each stimulus is associated with the amount and/or probability of reward. Retrospective calculation of stimulus-reward association cannot explain successful inferences based on traditional adjacent-pair training, because the frequency of reward is equal across all non-terminal stimuli. Nevertheless, more elaborate associative models have been proposed, such as value transfer theory (von Fersen et al., 1991), which posits that associative strength can “leak” between stimuli (such that stimulus A, being a “proven winner,” imparts some of its value to B). This, in principle, could permit TI-like effects in some scenarios without necessitating a full-blown representation of the ordered list.

Lazareva & Wasserman (2012) performed an important experiment to rigorously test several associative models of TI, including value transfer theory. They argued that previous studies of TI had guaranteed that all adjacent pairs received equal exposure during training. As a result, the empirical literature had not established whether human or animal performance would be disrupted if some pairs were presented more often than others. If, for example, the pair DE was presented much more frequently than the pair BC, then the selection of D would be paired with reward delivery more often than the selection of B. Value transfer theory, along with other associative models (e.g. Siemann & Delius, 1998), predicts that when presented with a subsequent pairing of BD, subjects should favor D over B as a result of D's greater association with reward. In general, presenting one pair repeatedly and reinforcing the choice of one item from that pair should drive that item's“reward value” toward ceiling.

With this in mind, Lazareva and Wasserman first trained pigeons on four adjacent pairs in a five-item list, presenting each pair with equal frequency. They then presented only the pair DE for hundreds of trials before they assessed TI performance. They also performed computer simulations to train and test whether associative models could perform TI following “massed trials” training to one pair. The results were clear: Pigeons who received massed DE trials showed equivalent performance to other animals who were trained equally on all pairs, and both groups consistently favored B over D. However, each associative model's performance was disrupted by massed presentation of DE, with performance either reduced to chance or displaying a heavy initial bias toward choosing D.

Lazareva and Wasserman's results make two things clear about the TI literature. First, the associative models they tested were conceived with a narrow set of task designs in mind. TI is ordinarily trained with equalized frequencies for all stimulus pairs, but this constraint shouldn't be expected in real-world choice scenarios. It is implausible that animals evolved with perfect counterbalancing as a feature of the environment, so models of transitive inference should be resistant to unequal presentation frequencies. Secondly, the development of more robust models of TI requires new experiments that manipulate previously uniform task parameters. By definition, although associative models seek to maximize positive outcomes and minimize negative ones, they only come to optimal conclusions when training carefully balances exposure to all outcomes. A key benefit of representation-based learning is that representations can decouple the relationship between stimuli and their individual reward histories, and this renders their inferences robust against biased sampling.

Although humans, rhesus monkeys, and pigeons all display serial learning, they do not do so in identical ways. Scarf & Colombo (2008) are blunt about the difference: “Simply put, birds are not as smart as primates” (p. 311). Subsequent studies of TI in birds have yielded TI performance comparable to rhesus monkeys (e.g. Bond et al., 2010), yet any similarity between cognitive functions in birds and mammals may be the result of wholly divergent brain architecture (Güntürkün & Bugnyar, 2016). Lazareva and Wasserman's demonstration that pigeon performance is not disrupted by massed trials has yet to be tested in primates.

It is also unclear from the literature how humans and monkeys might differ from one another under such a manipulation, because few comparative studies have pitted TI performance in humans against that of rhesus macaques directly (examples include Merritt & Terrace, 2011; Jensen et al., 2015). Most human studies are confounded by verbal instruction. Greene et al. (2001) compared the TI performance of human participants who either received explicit instructions, or who instead only learned by trial and error without verbal instructions. Although the latter group performed less well, both groups demonstrated above-chance performance on critical test pairs. Martin & Alsop (2004), however, provided a more nuanced result, splitting participants not in terms of whether they were given explicit instructions, but rather whether they displayed awareness of task demands during debriefing. Unaware participants did not significantly differ from chance on the critical test pairs.

We performed an experiment in which both rhesus macaques and human participants learned 7-item lists. In both cases, training began with massed presentation of the last two list items, pair FG. Since F was always correct when paired with G, the initial reward likelihood associated with F approached 100%. After a block of FG pairs, subjects then were trained on all pairs that included either F or G (to test what effect FG training had), as well as the adjacent pairs AB, BC, CD, and DE. Then, in a final phase, all 21 possible pairs were tested. If association with reward was a substantial influence, then subjects should develop a bias toward choosing F. Such a bias should disrupt performance during the second phase of the experiment, subsequent learning during that phase, or both.

## 2 Methods and Materials

### 2.1 Animal Subjects

Data were collected from 3 adult male rhesus macaques (*Macaca mulatta*): F, H, and L. All subjects had prior experience performing transitive inference tasks, as described by Jensen et al. (2015). Subjects were housed at the New York Psychiatric institute and its Department of Comparative Medicine, under the oversight of Columbia University's Institutional Animal Care and Use Committee (protocol AAAN7101). All operations were in accordance with the recommendations of the Guide for the Care and Use of Laboratory Animals of the National Institute of Health.

### 2.2 Animal Apparatus

Subjects were seated in an upright primate chair during the experiment. Head movements were restrained by head post, and eye movements were recorded using a monocular scleral search coil (Robinson, 1963; Judge et al., 1980) (CNC Engineering, Seattle WA). Eye movements (saccades, followed by fixation) were used by the animal to signal which stimulus was selected during each trial. Unless otherwise indicated, the apparatus was identical to that used by Teichert & Ferrera (2014).

### 2.3 Human Participants

Participants were 33 college undergraduates who earned course credit. The experiment was approved by Columbia University's Institutional Review Board (protocol AAAA7861), confirming to the guidelines for human research set forth by the American Psychological Association.

### 2.4 Human Apparatus

Participants performed the experiment on a personal computer (iMac MB953LL/A), with responses made via a mouse-and-cursor interface.

### 2.5 Procedure

At the start of the session, the task specified a list order for a set of seven stimuli, here denoted by the letters A through G (Figure 1 top). Subjects never observed all seven stimuli simultaneously; instead, each trial displayed a pair of list items. When stimuli were presented, a response was considered“correct” if it was made to the earlier item in this list. The structure of each trial is depicted in Figure 1 (left): Following a response to a start stimulus, subjects were then presented with the list pairing. Feedback regarding whether the response was correct (green check mark) or incorrect (red X) was given immediately. Animal subjects also received water via a tube for each correct response.

An experimental session always consisted of a fixed number of trials. Animal subjects completed 760 trials in a session, arranged into three phases (Figure 1 right). During Phase 1, the pair FG was presented 40 times. During Phase 2, 15 of the possible pairings were presented: All “adjacent pairs” (those having a symbolic distance of 1), and all other pairs that included either F or G (regardless of symbolic distance). The set of all pairings of F with a novel stimulus are hereafter denoted as “xF,” and all novel pairings with G are denoted as “xG.” The aim of this phase was to train the ordering of stimuli A through E relative to F and G, as well as to test performance on F-and-G-related pairs. Each of the 15 pairs was presented 20 times, resulting in a total of 300 trials. Finally, in Phase 3, all 21 pairings were presented. Again, each pair was presented 20 times, so Phase 3 lasted 420 trials. In all phases, the positions of the stimuli were randomly counterbalanced, such that the correct response was on each side of the screen half the time. Each subject completed multiple sessions (34 sessions for F, 76 sessions for H, and 20 sessions for L), with a new set of stimuli learned during each of those sessions.

**Figure 1:**
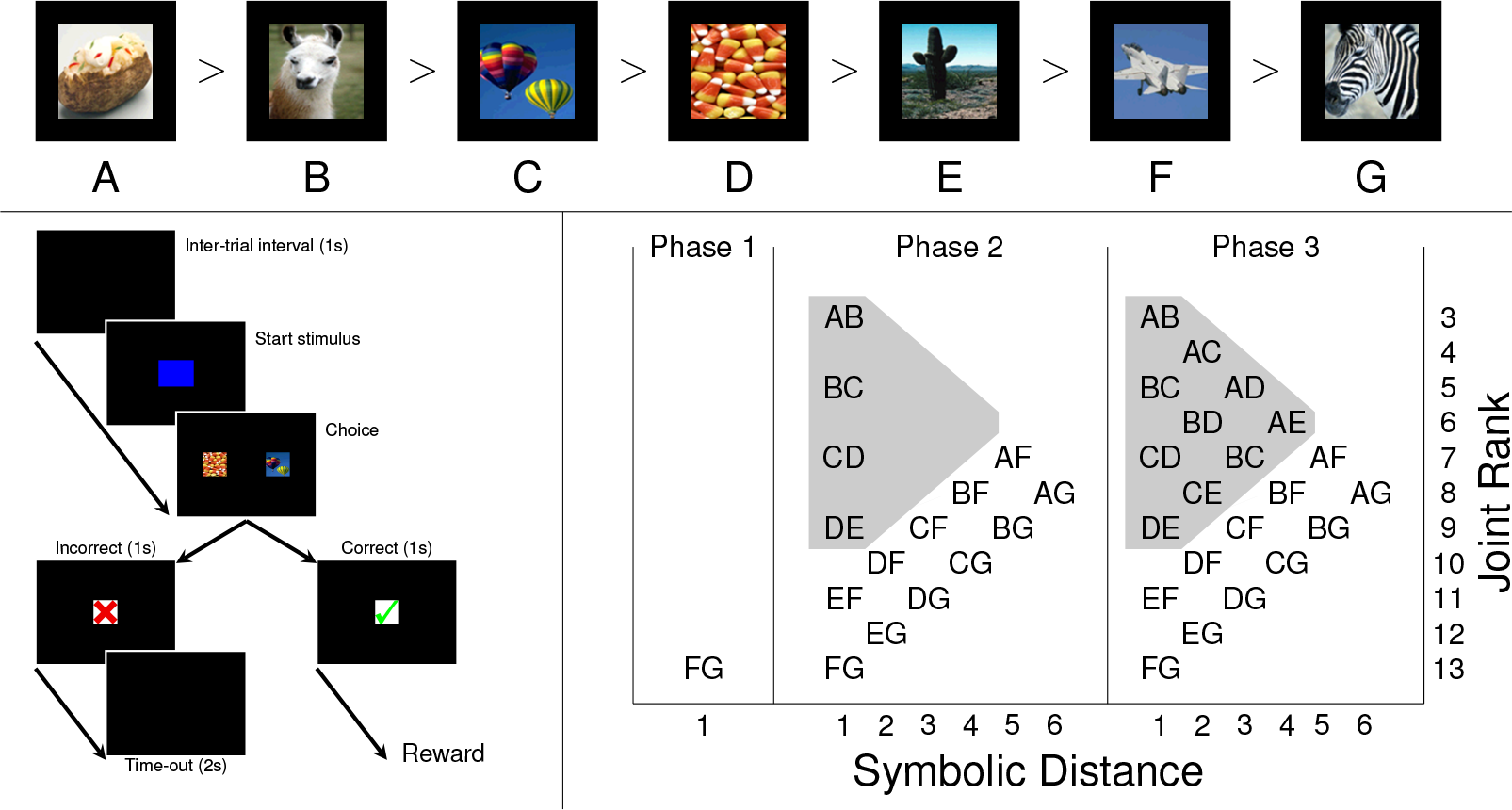
**Task and experimental design. Top Row.** Example of the stimuli in a 7-item transitive inference list. Since B > C > D, C is a correct answer when paired with D (or any stimulus of lower rank), but is incorrect when paired with B (or any stimulus of higher rank). **Bottom Left**. Sequence of events in a single trial. After an inter-trial interval, a blue start stimulus appears. Selecting it brings up a pair of images from the implied list. If the correct item is chosen, a green check mark is displayed, followed by reward delivery. If the incorrect item is chosen, a red X is displayed, followed by a 2 second time-out. **Bottom Right**. Design of experimental phases. In Phase 1, subjects are presented with only FG; in Phase 2, all adjacent pairs, as well as all pairs including F or G, are trained; in Phase 3, all pairs are trained. Pairs within the gray triangle were all stimuli that excluded F and G.

Figure 1 (right) shows how all 21 pairs are organized with respect to two predictors. The first, symbolic distance, is well-represented in the literature: It is the absolute difference between stimulus ranks. The second is “joint rank,” which is the sum of the stimulus ranks (here, low ranks correspond to earlier list items). Thus, the pair BD would have a distance of 2 and a joint rank of 6. There are several motivations for describing pairs in terms joint rank. First, each stimulus pair corresponds uniquely with a particular combination of distance and joint rank (e.g. no pair other than BD has a distance of 2 and a joint rank of 6). This gives each pair a specific coordinate within the predictor space. Secondly, joint rank is strictly orthogonal to distance. The largest joint rank (FG, joint rank of 13) and the smallest (AB, joint rank of 3) both have a symbolic distance of 1. This permits distance and joint rank to be used simultaneously as predictors without a collinearity confound. The intuitive interpretation of these two descriptors is that distance corresponds to the relative dispersion of stimuli along the number line, whereas joint rank corresponds to their absolute position within the list.

Human participants experienced the same procedure, albeit with half as many trials in each condition. Thus, they experienced Phase 1 for 20 trials, Phase 2 for 150 trials, and Phase 3 for 210 trials. All learning occurred in one session.

## 3 Results

In order to evaluate how estimated performance changed as a function of learning, we used Gaussian process regression (Rasmussen & Williams, 2006). Gaussian process regression is a highly flexible non-linear estimation approach that relies on discovering the extent to which every observation co-varies with every other, given some prior metric for comparing the distance between observations. Thus, while observations that are close to one another influence one another more, they also influence one another as a function of their similarity (Lucas et al., 2015). This permits robust time-series analysis to be performed without constraining the data to a particular functional form. Although a full Gaussian process model can be computationally prohibitive to fit (requiring runtime O(n^3^) for *n* observations), we took advantage of approximation techniques implemented in the GPstuff toolbox (Vanhatalo et al., 2013) to greatly accelerate estimation.

We first estimated how response accuracy and reaction time changed over the course of the experiment, doing so independently for each stimulus pair in each phase. Responses, coded as correct (1) or incorrect (0), were modeled using a binomial prior to obtain a continuous probability of correct responses. Reaction times were presumed to be approximately Gaussian on a log scale.

**Figure 2:**
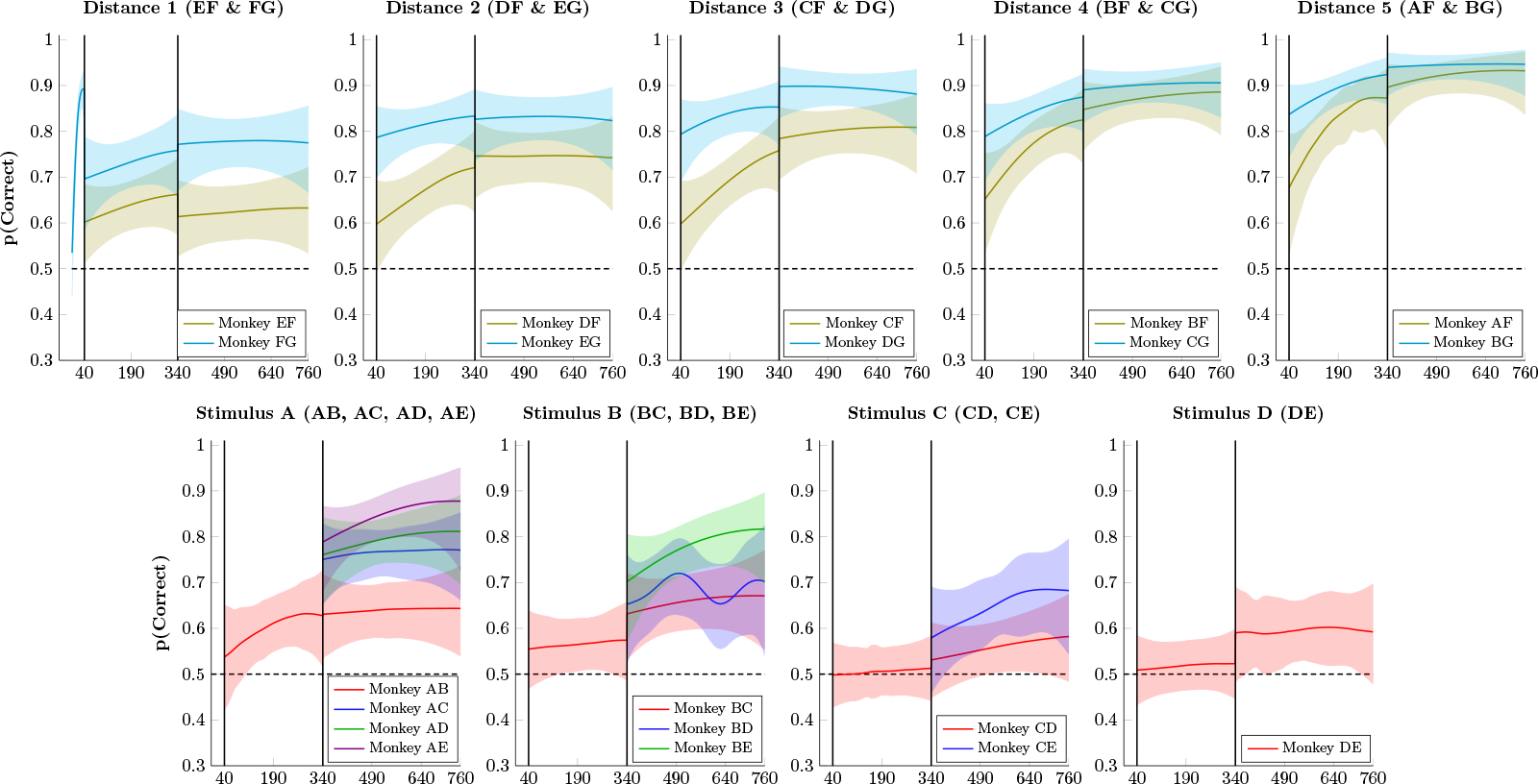
**Proportion of correct responses by monkeys for stimulus pairs over successive trials**. Shaded areas represent 99% credible intervals. **Top Row**.Performance on stimulus pairs that included F and/or G for distances 1 through 5. Performance on all such pairs exceeded chance levels, and performance on xG pairs consistently exceeded that of xF pairs. Despite rapidly climbing to nearly 89.0% accuracy, the pair FG dropped immediately to 69.6% accuracy at the start of Phase 2. **Bottom Row**. Performance on pairs composed of items A through E. Performance on adjacent pairs generally remained low during Phase 2, but improved somewhat in Phase 3. Additionally, the six novel pairs in Phase 3 showed enhanced performance, consistent with a symbolic distance effect.

Figure 2 (top row) shows mean response accuracy for monkeys choosing pairs that included either stimulus F (olive) or stimulus G (cyan). During the 40 trials of phase 1 (presentation of FG only), response accuracy on the pair FG rapidly improved, reaching approximately 90% correct selection of F. At the start of phase 2, however, accuracy for FG dropped to 70% accuracy. Phase 2 also began with above-chance performance for all pairs that paired either F or G with a novel stimulus (A thru E). This is unsurprising in the case of xG pairings because choosing G did not yield a reward in any phase of the task. However, consistent above-chance responding on xF pairs during phase 2 entails *avoidance* of F, despite F having yielded reliable rewards during phase 1. For example, the first time EF was presented (Fig. 2, top right, trial 41), E was chosen 60% of the time despite the fact that it had never been previously reinforced or even presented to the subject. Similar results were obtained for all of the novel stimuli. Thus, prior preference for F while it was paired with G did not prevent subjects from rapidly learning to not choose it when paired with the novel stimuli A through E.

**Figure 3:**
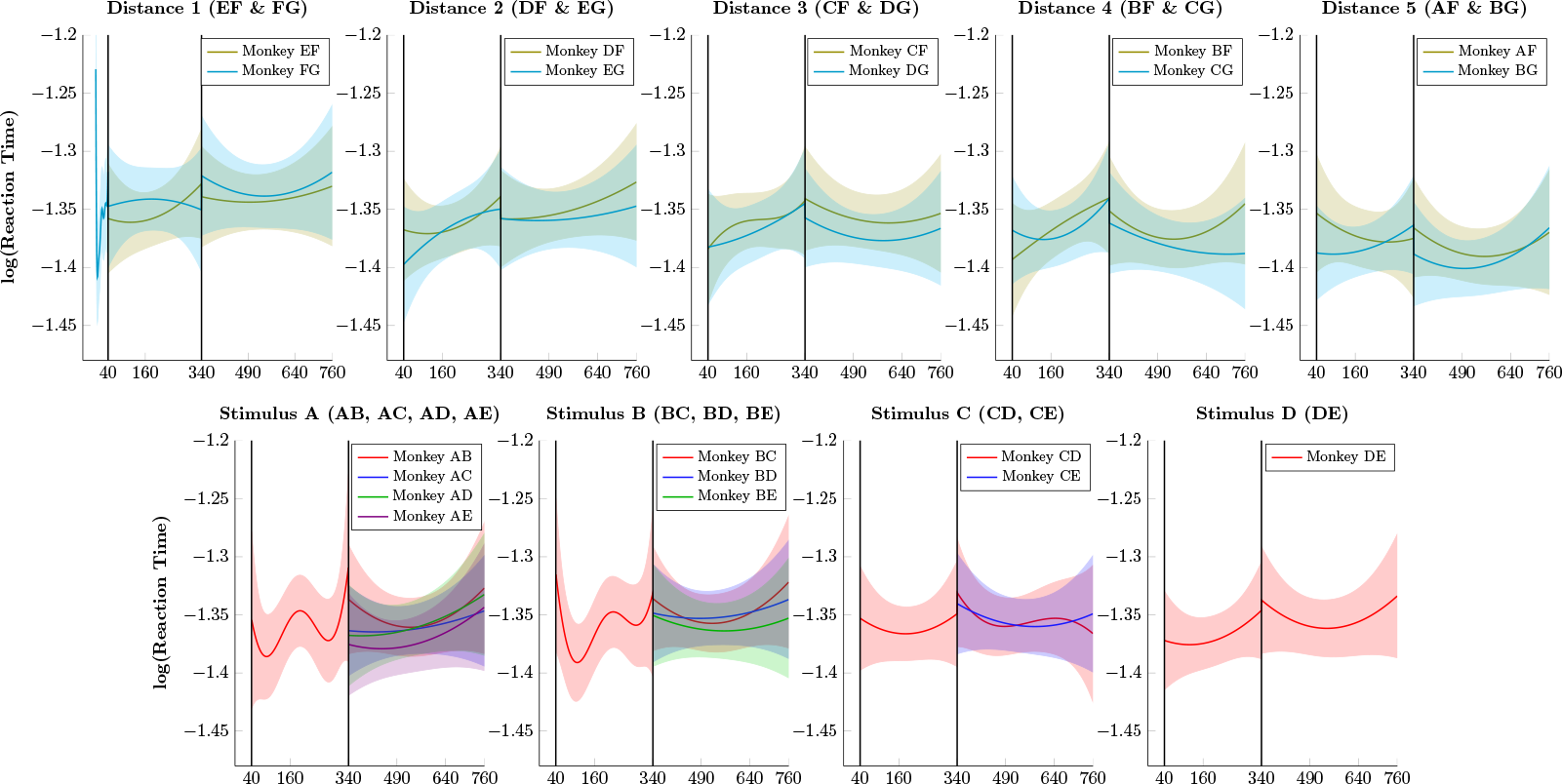
**Monkey reaction times (log-transformed) over successive trials**. Shaded areas represent 99% credible intervals. **Top Row**. Log reaction times for pairs that included F and G for distance 1 through 5. Reaction times were generally stable, with slightly faster times observed at larger symbolic distances. **Bottom Row**. Log reaction times for pairs composed of items A through E. Consistent reaction times were observed across all such pairs.

In phase 3, all 21 stimulus pairs were presented. By the end of phase 3, a very pronounced symbolic distance effect was evident: Accuracy for distance 1 pairs tended to be lowest, whereas the highest accuracy was associated with the distance 5 pairs. Figure 2 (bottom row) shows mean response accuracy for pairs composed of the first five items in the list, A through E. These are color-coded by symbolic distance (red = distance 1, blue = distance 2, green = distance 3, violet = distance 4). Adjacent pair performance rose above chance in the case of AB, but remained approximately at chance levels during phase 2 for the other pairs. However, despite low performance, a symbolic distance effect was observed at the transfer to all pairs in phase 3.

Figure 3 presents the log reaction times of monkeys, following the same format as Figure 2. Monkeys tended to respond very quickly (exp(−1.35) ≈0.26s), as is typical for oculomotor choice reaction times. Although some very mild differences seemed evident (the pair AF evoked faster responses than the pair EF), it seems reasonable to conclude that reaction times were near a lower floor for the fastest times that subjects could in principle achieve.

**Figure 4:**
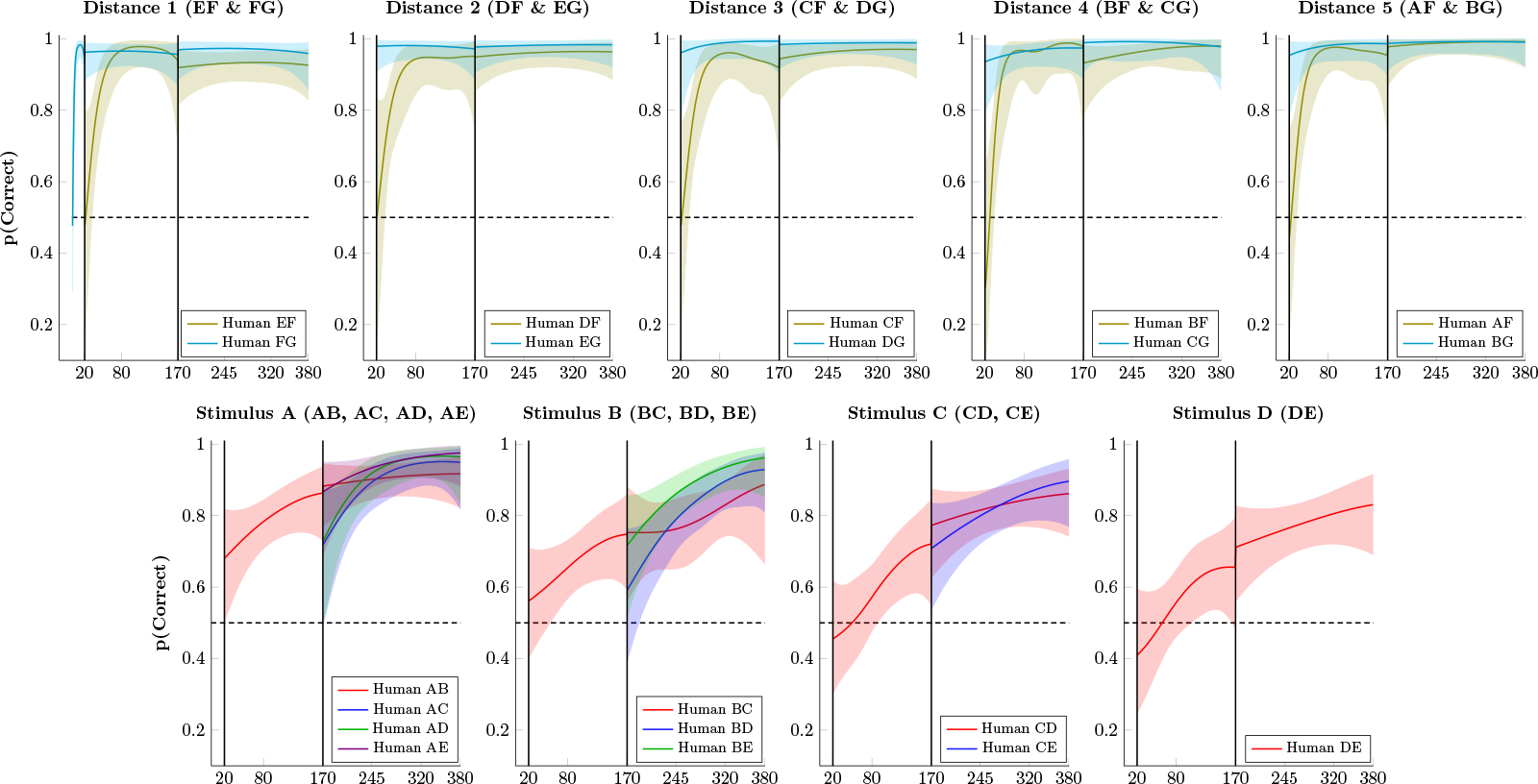
**Proportion of correct responses by humans for stimulus pairs over successive trials**. Shaded areas represent 99% credible intervals. **Top Row**. Performance on stimulus pairs that included F and/or G for distances 1 through 5. Performance on FG rapidly rose to ceiling levels, as did performance on pairs including G. Other pairs including F began at chance level at the start of Phase 2, but also rapidly rose to ceiling. **Bottom Row**. Performance on pairs composed of items A through E. Accuracy on adjacent pairs rose steadily on Phases 2 and 3, and novel non-adjacent pairs were above chance (but below adjacent pairs) at the start of Phase 3.

Figure 4 presents mean response accuracy for human participants. Unlike monkeys, participants acquired a stronger preference for F during phase 1. Despite this, they immediately discounted F at the beginning of phase 2. They then showed rapid acquisition, racing to ceiling on all xF and xG pairs. Performance also rapidly rose above chance levels for adjacent pairs in phase 2, and a symbolic distance effect was not evident. If anything, performance on adjacent pairs tended to slightly exceed the distance 2 pairs.

Figure 5 presents the log reaction times of participants. Humans responded much more slowly than monkeys (exp(0.5) ≈ 1.65s), but unlike monkeys they became systematically faster over the course of the session. Signs of a symbolic distance effect were more evident in the human reaction times than in their response accuracy. In particular, participants tended to respond more quickly to larger symbolic distances.

**Figure 5:**
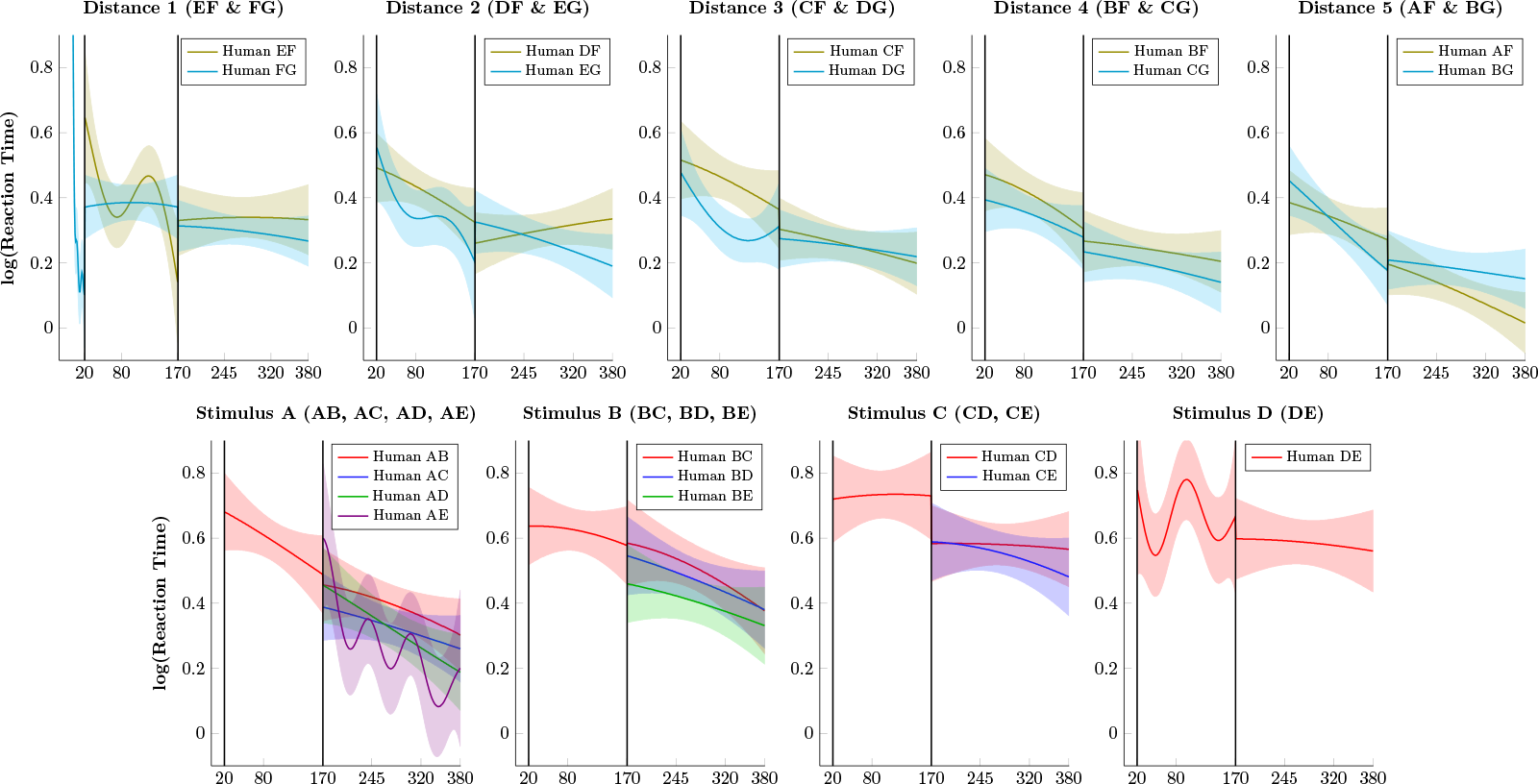
**Human reaction times (log-transformed) over successive trials**. Shaded areas represent 99% credible interval. **Top Row**. Log reaction times for pairs that included F and G for distance 1 through 5. Participants were much slower than monkeys, but reaction times steadily improved over the course of the experiment. **Bottom Row**. Log reaction times for pairs composed of items A through E. Reaction times to pairs that included item A sped up most during learning, whereas pairs closer to the “middle” of the list (such as DE) sped up very little.

The transition at the start of phase 2 shows several surprising effects with respect to F. At the start of phase 2, monkeys appeared to consistently avoid F, despite consistent reinforcement of its selection in phase 1. In addition, monkey response accuracy on FG appeared to drop abruptly on a single trial, from near 90% at the end of phase 1 to near 70% at the start of phase 2. Humans showed neither of these effects. Their FG trials remained close to ceiling, whereas all other xF pairs appeared to begin near chance (albeit with considerable uncertainty). The patterns of responding to F suggest different processes at work in these two species.

To get a more precise understanding of this transition, we examined choices based on order of appearance at this transition, as shown in Figure 6. The first four points report the estimated response accuracy for FG during the last four trials of phase 1 for both monkeys (Figure 6 left) and humans (Figure 6 right). The next six points in each plot report the nth instance in phase 2 of FG (black), any xF pair (olive), or any xG pair (cyan). For example, the first xF pair in phase 2 might be the pair AF or BF, and might not take place until the third or fourth trial. Thus, the first xF trial in phase 2 is the first unambiguous case where some other stimulus should be favored over F.

**Figure 6:**
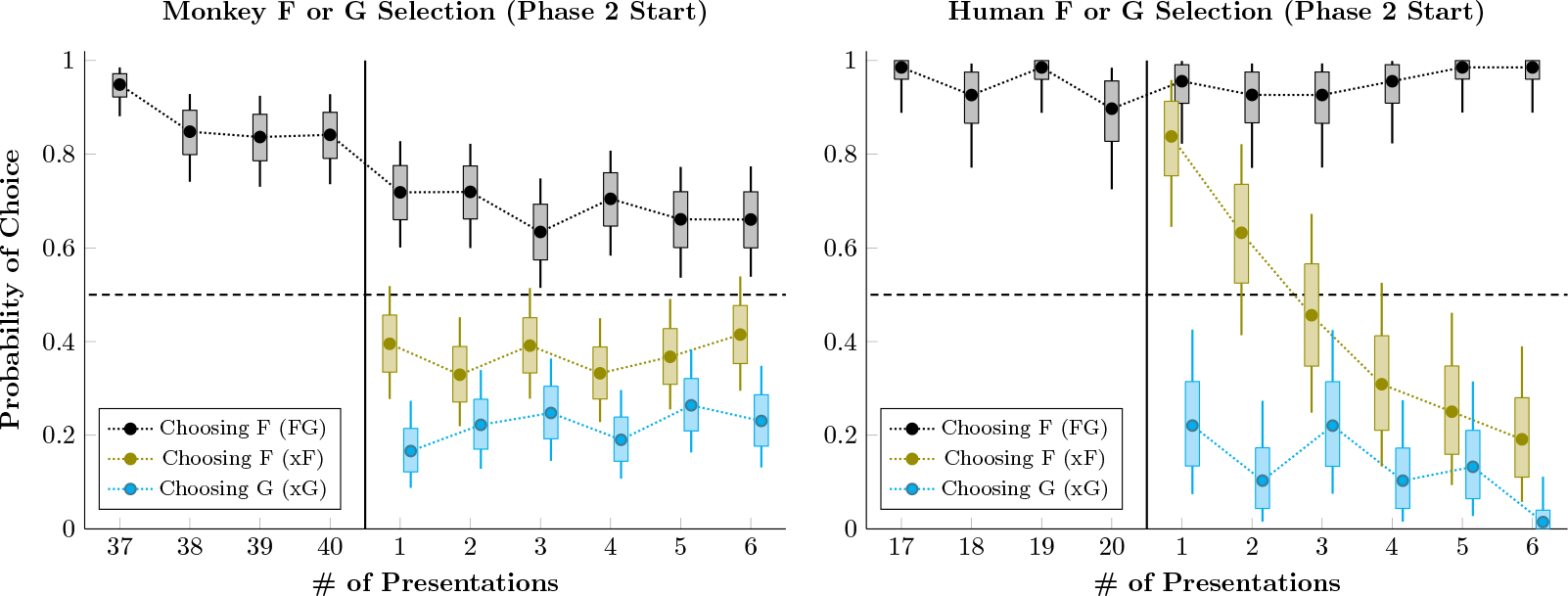
**Probability of choosing F and G during a pair's last four presentations during phase 1 and first six presentations during phase 2**. Whiskers represent the 99% credible interval for the parameter, and boxes represent the 80% credible interval. The probability of choosing F in the context of the pair FG (black) is contrasted against the probability of choosing F in the context of all other xF pairs (olive). Also shown is the probability of choosing G in the context of all other xG pairs (cyan). **Left**. Mean selection probability for rhesus macaques. Monkeys show a drop in performance between the last presentation of FG in phase 1 and its first presentation in phase 2. Additionally, monkeys avoid choosing F even on the first xF pairing. **Right**. Mean selection probability for human participants. Although humans avoid G in xG pairings, they show a brief preference for F during the first presentation of xF in phase 2. This preference has been fully reversed after only four presentations.

Figure 6 (left) shows that monkeys chose F less than half the time on the very first xF trial. It also shows that the drop in FG performance between phases is indeed an abrupt step downwards. Contrastingly, humans were about 80% likely to choose F during the first xF pairing, and about 20% likely to choose G during the first xG pairing. This preference for F was very rapidly corrected. Thus, whereas monkeys had a prior expectation that F should be avoided in favor of novel stimuli, humans had to instead rapidly “unlearn” the associated value of F.

Gaussian process regression is a continuous time-series analysis that permits estimates of not only response accuracy, but also the rate at which performance changes. Figure 7 shows the rate at which monkeys improved at the start of phases 2 and 3, the equivalent of the slope in Figure 2 measured with respect to log-odds. At the start of phase 2 (Figure 7 left), monkeys displayed an apparent symbolic distance effect with respect to the rate of learning, an effect not previously reported in the literature. In addition, learning was generally faster for xF pairs than it was for xG pairs, or for the entirely novel pairs (AB, BC, DC, DE). By the start of phase 3 (Figure 7 right), however, learning rates were consistently slow. Only novel pairs had a learning rate excluded zero from the 99% credible interval.

**Figure 7:**
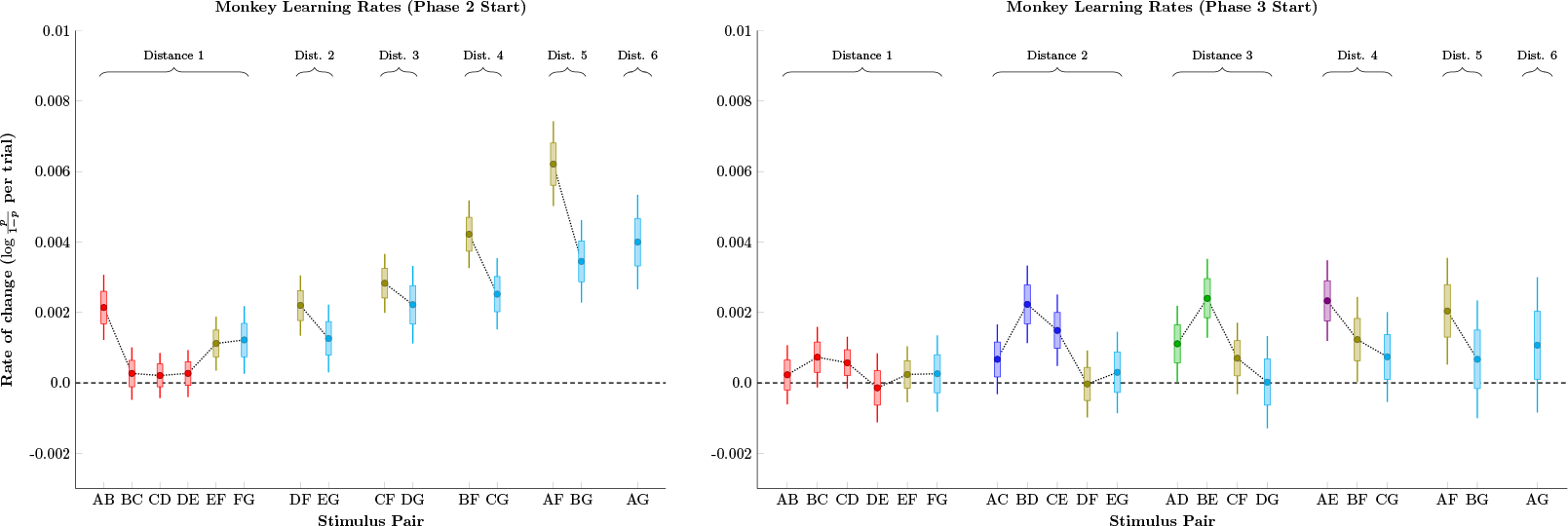
**Estimates of the rate of change of response accuracy for monkeys, measured in log-odds units of probability per trial**. Pairs are sorted by symbolic distance and by joint rank. Boxes represent 80% credible intervals, while whiskers represent 99% credible intervals. Individual pairs are color-coded identically to Figures 2-5. **Left**. Learning rate at the start of phase 2. Pairs including stimulus F tended to show faster improvement, as did pairs with greater symbolic distance. **Right**. Learning rate at the start of phase 3. Novels pairs showed elevated learning, but all learning rates were low compared to phase 2.

Figure 8 shows the corresponding learning rate analysis for human participants, which were very different than those of monkeys. First, human learning rates were about an order of magnitude faster. Secondly, no symbolic distance effect was evident. Instead, xF pairs were learned very rapidly at the start of phase 2 (consistent with Figure 6), and had stabilized by the start of phase 3. Another difference was that novel pairs in phase 3 seemed to be learned faster when paired with stimulus close to A. This is likely an example of a terminal item effect, which is only seen at one end of the list because the other end has already received considerable training.

To better understand how the symbolic distance effect and the terminal item effect might manifest in these data, a more complex Gaussian process model combined several predictors. This permitted estimates to be fit that combined all stimulus pairs. Trial number was used as a predictor, in order to make a precise inference of performance during the first trial of Phase 3. Additionally, each pair was coded for its symbolic distance and its joint rank. As noted above, the orthogonality of distance and joint rank permits the two to be used simultaneously as predictors across all pairs. Additionally, estimates may also be obtained for the interpolated regions between stimuli, even though no stimulus pairing resides in those regions. This facilitates understanding of the underlying estimated function.

**Figure 8:**
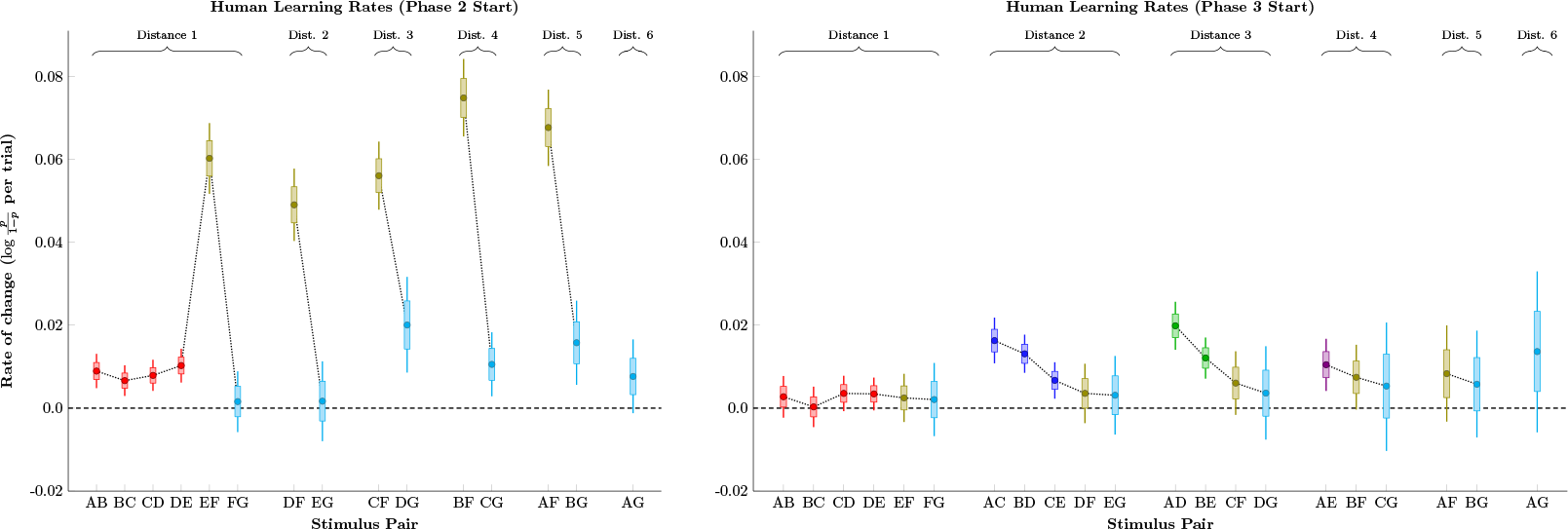
**Estimates of the rate of change of response accuracy for humans, measured in log-odds units of probability per trial**. Pairs are sorted by symbolic distance and by joint rank. Boxes represent 80% credible intervals, while whiskers represent 99% credible intervals. Individual pairs are color-coded identically to Figure 2-5. **Left**. Learning rate at the start of phase 2. Pairs including stimulus F tended to show much faster improvement. **Right**. Learning rate at the start of phase 3. Novels pairs showed elevated learning, and most learning rates were higher than those observed in monkeys.

Figure 9 shows the estimated probability of a correct response (top row) and the log reaction time (bottom row) for all monkeys at the start of Phase 3. The color coding of each of the pairs is consistent with that used in Figures 2 through 5, and the novel pairs at this point in the experiment are AC, BD, CE, AD, BE, and AE. Both response accuracy and reaction time show a consistent distance effect in both familiar and unfamiliar pairs, with larger symbolic distances associated with higher accuracy and lower reaction times. The added benefit of previously training (in phase 2) the pairs including F and G is evident in the asymmetry of these curves. The case for a terminal item effect in the monkey data are mixed, however. While pairs including the terminal A and G appeared to yield slightly higher accuracy, log reaction times seemed closer to linear, with FG being the slowest pair despite being the most extensively trained.

**Figure 9:**
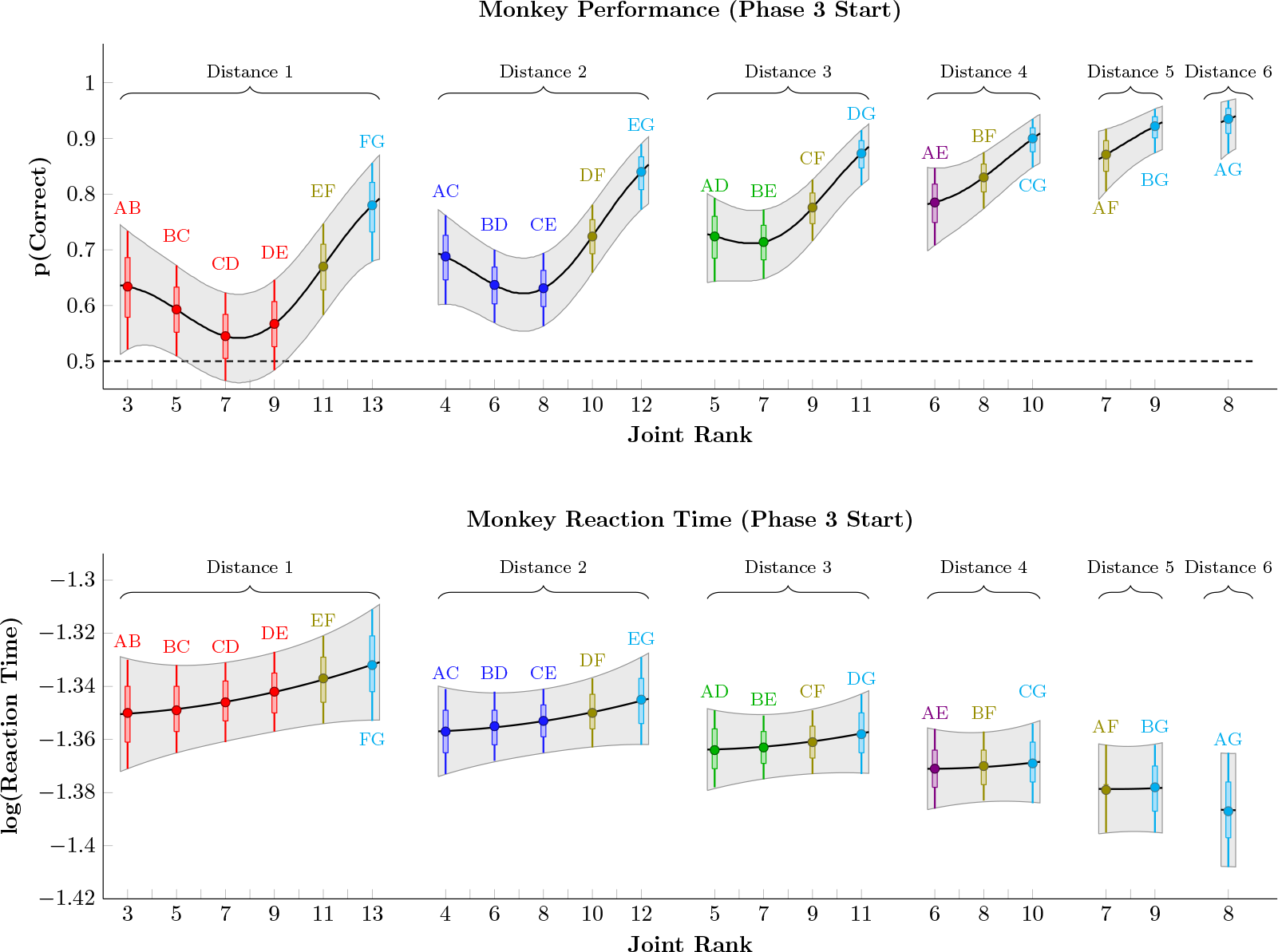
**Estimates of response accuracy (top row) and log reaction times (bottom row) in monkeys during the first trial of phase 3**. Pairs are sorted by symbolic distance and by joint rank. Gray shaded areas correspond to the 99% credible interval, interpolated between items. Individual pairs are color-coded identically to Figure 2-5, and also include a horizontal bar to denote 80% credible intervals. Response accuracy displayed both a symbolic distance effect and a terminal item effect, as well as asymmetry resulting from the additional training on stimuli F and G. Log reaction times displayed a symbolic distance effect, but did not show a clear terminal item effect.

Figure 10 depicts response accuracy and reaction times for human participants at the start of Phase 3. As in Figures 4 and 5, accuracy was consistently higher, and reaction times slower, than in monkeys. Additionally, however, a much clearer pattern of terminal item effects was evident (especially for the pair AB). Although a clear distance effect was evident in the reaction times, the evidence for such an effect was much more equivocal in their accuracy. As in Figure 4, performance on adjacent pairs tended to marginally exceed that of corresponding distance 2 pairs. However, Distance 4 pairs tended to outperform distance 3 pairs, and distance 3 pairs tended to outperform distance 2 pairs, suggesting a mild distance effect among the six novel pairs. The effect of massed training of FG was also evident in human reaction times, such that xF and xG reaction times were consistently faster than other stimulus pairs.

## 4 Discussion

Learning and decision-making are thought to depend on an ability to predict the reward value of alternative actions. Reinforcement-based theories critically depend on the history of experienced rewards, and seek to explain choices made by non-human animals in terms of associations between stimuli, actions, and outcomes. Alternatively, cognitive factors appear to also contribute to animal decision making, whereby representational mechanisms both facilitate and constrain an organism's ability to infer relationships between stimuli and outcomes. The transitive inference paradigm (TI) can help disambiguate these contributions, because explaining TI performance in terms of stimulus-reward associations is especially difficult. Our results instead support cognitive interpretations of TI performance.

Adapting a manipulation introduced by Lazareva & Wasserman (2012), we investigated whether TI performance was disrupted by additional training on a particular stimulus pair. According to associative models, when the pair FG is overtrained, it should drive the association of stimulus F with reward toward ceiling. If TI depends on stimulus-reward associations, the reward history of F should lead to it being favored in pairings such as EF and DF. Even though we repeatedly presented FG trials in advance of ordinary TI learning in both rhesus monkeys and humans, we found no evidence of a bias for choosing F in subsequent phases of learning. This result is consistent with the lack of disrupted learning in pigeons reported by Lazareva and Wasserman.

**Figure 10:**
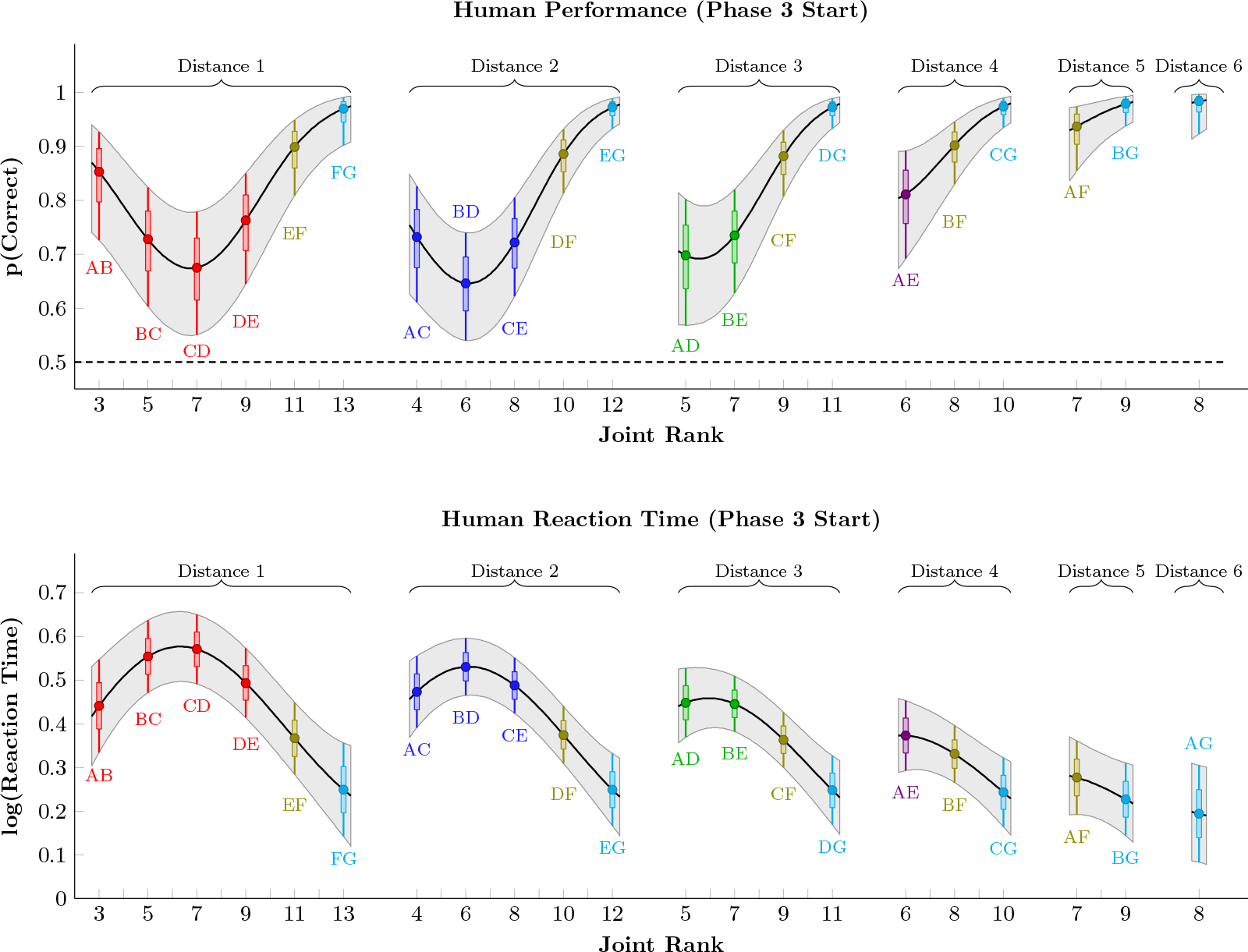
**Estimates of response accuracy (top row) and log reaction times (bottom row) in humans during the first trial of phase 3**. Pairs are sorted by symbolic distance and by joint rank. Gray shaded areas correspond to the 99% credible interval, interpolated between items. Individual pairs are color-coded identically to Figure 2-5, and also include a horizontal bar to denote 80% credible intervals. Human response accuracy showed a pronounced terminal item effect, but a weak symbolic distance effect. Human reaction times, on the other hand, displayed both effects.

Following FG training, rhesus macaques displayed a surprising pattern of behavior. Although novel stimuli paired with G were more likely to be selected than novel stimuli paired with F, all xF and xG pairings yielded performance above chance. In other words, despite a learning history in which F had only ever yielded reward, subjects systematically avoided F in favor of novel stimuli. This result, evident in Gaussian process estimates of response accuracy (Figure 2), was also found in the accuracy of the very first xF trial of phase 2 (Figure 6). Every session was performed with novel stimuli, so subjects could not have had a prior belief about any specific stimulus in the F position. However, because subjects learned many lists over the course of the experiment, it is likely that this above-chance performance was the result of a “task representation” (Stoet & Snyder, 2003), sometimes called a “task set” (Sakai, 2008). Despite having never seen stimulus F before the current session, subjects had learned that stimuli like F (i.e. those sharing the characteristic of having appeared during the initial massed trials) were likely to be of lower rank than novel stimuli seen later in the session. The strong preference for F established early in the session would thus be conditional, and F would immediately be avoided when paired with novel stimuli.

Another surprising effect was that despite achieving approximately 90% accuracy overall during FG training, accuracy on the pair dropped to around 70% as soon as other stimulus pairs began to be presented. This effect was evident even when comparing, on a session-by-session basis, the last FG trial of phase 1 to the first FG trial of phase 2. While this could potentially be another manifestation of a task representation, another possibility is that the maintenance of a stimulus pair and inferential reasoning both draw upon the same working memory resources (Halford et al., 2007). This is consistent with past studies that have found that rhesus monkeys perform only slightly above chance on adjacent pairs after hundreds of training trials (Jensen et al., 2013, 2015). Despite only needing to learn the correct response for 6 adjacent pairs, monkeys seemed unable to memorize the correct choice for each pair. Past estimates of the capacity of visual short-term memory suggest that rhesus macaques are limited to between one and two items (Elmore et al., 2011). Thus, poor performance on pairings that animals have seen many times over may reflect the unavailability of memorization strategies, leaving more implicit mechanisms to perform the necessary inferences. The present result (wherein FG is selected with high accuracy only when presented in a block of identical trials) suggests that performance in phase 1 was facilitated by a memory trace of the previous trial.

Our human participants displayed a different pattern of behavior from the monkeys. Superficially, this pattern more closely resembled the expectations from associative models: Participants favored F on the very first xF pairing, whereas they avoided G on the first xG pairing. Humans adapted very rapidly, however, correctly avoiding F by the fourth xF presentation. The speed of human adaptation to xF pairs was much faster than any other learning displayed in this study, which is especially notable given that participants were task-naive. Associative models with very fast learning rates are prone to unstable behavior, on account of their oversensitivity to strings of good and bad luck. Furthermore, the humans’ very rapid learning did not appear to be a feature of the transition from phase 2 to phase 3, as would be expected if rapid learning was a mere consequence of an associative learning rate.

In both the monkey and human cases, the stimuli A through E effectively offered a second, embedded TI task, with adjacent-pair training during phase 1 and all-pair testing during phase 3. Humans and monkeys both successfully inferred the relationships between nonadjacent test pairs from this set with no evidence of disruption due to overtraining of F.

Overall, the present human and monkey results display both consistencies and differences with Lazareva and Wasserman's pigeon data. On the one hand, FG training did not disrupt learning of stimuli A through E. Monkeys never favored F, despite initial rewards earned from FG. They appeared to exploit a task representation to anticipate F?s undesirability relative to novel stimuli. Beyond that anticipation, their performance otherwise displays classical symbolic distance and terminal item effects. One of the most curious effects was a drop in the accuracy of FG choice at the start of phase 2.

Human behavior at the start of phase 2 did not resemble that of monkeys. Participants very briefly favored F, but they overcame their initial bias almost immediately. They also did not display drops in response accuracy to any pairings following phase transitions.

### 4.1 Time Series Analysis of TI

Traditionally, studies of TI train subjects extensively on adjacent pairs, then “test” them using non-adjacent pairs. The reported duration of training in such studies is not consistent from one paper to the next. Many labs end training based on a performance criterion, so subjects may experience training periods of dramatically different lengths. Since the aim of a behavioral test is to assess performance during cross-section in time, all that can be ascertained during test trials is response accuracy and reaction time at the moment of the test.

In this study, we instead approached TI performance as a time series, with the expectation that subjects would continue to accrue information over the course of successive trials, and that performance would improve accordingly. Using Gaussian process regression, we were able to construct a much more detailed picture of how learning unfolded for each stimulus pair in both species.

Rather than report performance as a function of stimulus pair, Figure 7 depicts the rate of change for monkey response accuracy, doing so in terms of log-odds units of probability. As far as we are aware, the only method previously used to measure learning rate in studies of TI has been “trials to criterion” (e.g. Gazes et al., 2012) making ours the first study to examine the first derivative of response accuracy directly. Figure 7 reveals an intriguing possibility: A symbolic distance effect for learning rate. The implication is that, when measured in terms of log-odds, stimulus pairs separated by larger symbolic distances are not only expected to have higher accuracy, but also to improve more rapidly as additional training unfolds. Realistically, this effect can't last: As performance reaches ceiling, learning rates drop to zero. Nevertheless, there is no evidence that prior overtraining of F impaired the rate at which monkeys learned to avoid F.

Figure 8 depicts the rate of change for human performance. The difference in scale from monkey learning is striking, with the mean learning rate as much as ten times faster than that of monkeys. The pattern of learning was qualitatively different between the two species: At the start of phase 2, humans made steady progress on most pairs (including all novel adjacent pairs), and also made very rapid progress on all xF pairs. Although it is intuitively obvious that humans monkeys differ in their rate of learning (c.f. visual comparisons of performance in Figure 2 to Figure 4), the analysis of the learning rate provides a more quantitative account of that difference.

Comparisons of learning rates also provide a new way to compare the predictions of different models of TI. This, in turn, can be used to place much more severe constraints on models that seek to explain behavior. Additional studies will be needed to determine how learning rate effects manifest in different species and in various experimental preparations.

### 4.2 Task Awareness & Task Representation

Task “awareness” has been a major focus in the human study of TI (Greene et al., 2001; Martin & Alsop, 2004), and it has been proposed as an explanation of the difference between human and monkey TI performance (Moses et al., 2006). In most cases, task awareness is defined in terms of verbal report: If a participant can deduce (and subsequently report) that the stimuli belong to an ordered hierarchy, they are considered to be “aware;” all others are “unaware.” Those participants who could not verbally articulate the structure of the task also perform very poorly on the inferential test trials (Frank et al., 2005; C. Smith & Squire, 2005; Libben & Titone, 2008).

Unfortunately, the way awareness has been defined does little to clarify the underlying cognitive mechanisms of TI. Although it appears as though awareness and successful inference for test pairs are connected in humans (Libben & Titone, 2008; Titone, 2008), it is unclear whether the two can be dissociated and, if so, which precedes the other. Moses et al. (2008) argue that TI depends not on a single brain region or cognitive faculty, but rather on “cognitive integrity” across multiple subsystems.

While it is true that monkeys differ from “aware” humans in a variety of respects, they also differ from “unaware” humans in an important way: the monkeys display transitive inference, whereas unaware humans do not. This raises an important question: If task awareness is linked to TI in humans, are analogous mechanisms involved in non-human TI? The “awareness” label in the human literature relies on verbal report, so macaques by definition do not qualify. Instead, a range of theoretical constructs regarding animal cognition have been proposed whose functions may overlap with the task awareness reported in humans.

The observed above-chance performance on xF pairs, in which animals avoided F despite its reward history, suggests that monkeys employed a “task representation” (Stoet & Snyder, 2003; Sakai, 2008). It is unlikely that this effect was driven by the saliency of the novel stimuli with which F was paired, as other novel pairs (BC, CD, DE) were learned slowly. Task representations, in this context, operate as superordinate frameworks within which current learning is embedded. Thus, as subjects learn how to respond to the current session's pair FG, they are also presumably learning how F and G fit into a broader context learned during previous sessions.

The comparative study of task representations has yet to reach a consensus regarding either terminology or theoretical focus. The study of how animals learn overarching task demands has also been described as “rule representation” (Bunge & Wallis, 2008), “abstract concept learning” (Wright & Katz, 2007), or simply “task context” (Asaad et al., 2000). Although the evidence suggests representations more abstract than those proposed by associative models, the available evidence falls short of a demonstration of “task awareness,” which as defined in humans is explicitly metacognitive (Terrace & Son, 2009). Although several lines of behavioral evidence are consistent with hypothesized metacognitive faculties in animals (J. D. Smith et al., 2014), whether metacognition can be inferred from behavior alone may be undecidable (Clark & Hassert, 2013).

Consequently, building a bridge between task awareness and task representation will require a better understanding of the implicit mechanisms of human task representation. When the evidence is approached from this perspective, there are good reasons to think that humans differ from macaques with respect to how task representations influence behavior. Humans, for example, are adept at “task switching” paradigms. These require participants to make choices according to one of several possible rules (Kiesel et al., 2010). When faced with task switching, rhesus macaques have difficulty suppressing the inappropriate rule. Instead, both behavioral (Avdagic et al., 2000) and neurophysiological (Klaes et al., 2011) evidence suggest that macaques consider both rules simultaneously, resulting in interference. One pair of studies makes this species difference especially clear. Stoet & Snyder (2003) demonstrated that, over the course of tens of thousands of trials, macaques had difficulty using a cue to switch from one task representation to another. Humans performed an identical procedure for an equally lengthy training period, and never displayed interference between task sets (Stoet & Snyder, 2007).

In this spirit, future comparative studies of TI should place greater emphasis on exploring how task representations are constructed, maintained, and deployed by different species. The first step in doing so is to study how task representation contributes to TI performance in humans without falling back on self-report.

## 5 Conclusion

Initial overtraining that repeatedly associated a single stimulus with reward did not impair monkeys’ or humans’ ability to avoid choosing that stimulus when appropriate in the context of TI learning. Instead, both species used information gleaned from the task to improve their performance, albeit in different ways. Monkeys, having extensive experience with the task structure, were able to identify the stimuli F and G as belonging to a particular class of stimuli across sessions, which in turn let them begin phase 2 above chance. Humans, on the other hand, were task-naive, and showed an initial preference for F which was almost immediately replaced with avoidance. The difference between the species were further evidenced in a novel analysis of the learning rate. Although the relative association of stimuli with reward is a useful strategy under some circumstances, it does not appear to be how transitive inference is performed. Instead, cognitive models that relate stimuli to one another along some ordinal dimension, as well as represent general task structure, are needed to account for the full range of published TI results.

## 6 Acknowledgments

The authors wish to thank Carolina Montes, Orly Morgan, Grant Spencer, and Nathan Tumazi for their assistance collecting the human data.

## References

Allen, C. (2006). Transitive inference in animals: Reasoning of conditioned associations. In S. Hurley & M. Nudds (Eds.), Rational animals? (pp. 175–185). London, UK: Oxford University Press.

Asaad, W. F., Rainer, G., & Miller, E. K. (2000). Task-specific neural activity in primate prefrontal cortex. Journal of Neurophysiology, 84, 451–459.

Avdagic, E., Jensen, G., Altschul, D., & Terrace, H. S. (2000). Rapid cognitive flexibility of rhesus macaques performing psychophysical task-switching. Animal Cognition, 17, 619–631.

Bond, A. B., Wei, C. A., & Kamil, A. C. (2010). Cognitive representation in transitive inference: A comparison of four corvid species. Behavioural Processes, 85, 283–292.

Bunge, S. A., & Wallis, J. D. (Eds.). (2008). Neuroscience of rule-guided behavior. New York, NY: Oxford University Press.

Clark, K. B., & Hassert, D. L. (2013). Undecidability and opacity of metacognition in animals and humans. Frontiers in Psychology, 4, Article 171.

Couvillon, P. A., & Bitterman, M. E. (1992). A conventional conditioning analysis of “transitive inference” in pigeons. Journal of Experimental Psychology: Animal Behavior Processes, 18, 308–310.

D'Amato, M. R., & Colombo, M. (1990). The symbolic distance effect in monkeys (cebus apella). Animal Learning & Behavior, 18, 133–140.

Elmore, L. C., Ma, W. J., Magnotti, J. F., Leising, K. J., Passaro, A. D., Katz, J. S., & Wright, A. A. (2011). Visual short-term memory compared in rhesus monkeys and humans. Current Biology, 21, 975–979.

Frank, M. J., Rudy, J. W., Levy, W. B., & O'Reilly, R. C. (2005). When logic fails: Implicit transitive inference in humans. Memory & Cognition, 33, 742–750.

Gazes, R. P., Chee, N. W., & Hampton, R. R. (2012). Cognitive mechanisms for transitive inference performance in rhesus monkeys: Measuring the influence of associative strength on inferred order. Journal of Experimental Psychology: Animal Behavior Processes, 38, 331–345.

Greene, A. J., Spellman, B. A., Dusek, J. A., Eichenbaum, H. B., & Levy, W. B. (2001). Relational learning with and without awareness: Transitive inference using nonverbal stimuli in humans. Memory & Cognition, 29, 893–902.

Güntürkün, O., & Bugnyar, T. (2016). Cognition without cortex. Trends in Cognitive Sciences, 20, 291–303.

Halford, G. S., Cowan, N., & Andrews, G. (2007). Separating cognitive capacity from knowledge: A new hypothesis. Trends in Cognitive Sciences, 11, 236–242.

Jensen, G., Altschul, D., Danly, E., & Terrace, H. S. (2013). Transfer of a serial representation between two distinct tasks by rhesus macaques. PLOS ONE, 8, e70285.

Jensen, G., noz, F. M., Alkan, Y., Ferrera, V. P., & Terrace, H. S. (2015). Implicit value updating explains transitive inference performance: The betasort model. PLOS Computational Biology, 11, e1004523.

Judge, S. J., Richmond, B. J., & Chu, F. C. (1980). Implantation of magnetic search coils for measurement of eye position: An improved method. Vision Research, 20, 535–538.

Kiesel, A., Steinhauser, M., Wendt, M., Falkenstein, M., Jost, K., Philipp, A. M., & Koch, I. (2010). Control and interference in task switching - a review. Psychological Bulletin, 136, 849–548.

Klaes, C., Westendorff, S., Chakrabarti, S., & Gail, A. (2011). Choosing goals, not rules: Deciding among rule-based action plans. Neuron, 70, 536–548.

Lazareva, O. F., & Wasserman, E. A. (2012). Transitive inference in pigeons: Measuring the associative values of stimuli b and d. Behavioural Processes, 89, 244–255.

Libben, M., & Titone, D. (2008). The role of awareness and working memory in human transitive inference. Behavioural Processes, 77, 43–54.

Lucas, C. G., Griffiths, T. L., Williams, J. J., & Kalish, M. L. (2015). A rational model of function learning. Psychonomic Bulletin & Review, 22, 1193–1215.

Martin, N., & Alsop, B. (2004). Transitive inference and awareness in humans. Behavioural Processes, 67, 157–165.

McGonigle, B. O., & Chalmers, M. (1977). Are monkeys logical? Nature, 267, 694–696.

McGonigle, B. O., & Chalmers, M. (1992). Monkeys are rational!. Quarterly Journal of Experimental Psychology, 45B, 189–228.

Merritt, D. J., & Terrace, H. S. (2011). Mechanisms of inferential order judgments in humans (homo sapiens) and rhesus monkeys (macaca mulatta). Journal of Comparative Psychology, 125, 227–238.

Moses, S. N., Villate, C., Binns, M. A., Davidson, P. S. R., & Ryan, J. D. (2008). Cognitive integrity predicts transitive inference performance bias and success. Neuropsychologia, 46, 1314–1325.

Moses, S. N., Villate, C., & Ryan, J. D. (2006). An investigation of learning strategy supporting transitive inference performance in humans compared to other species. Neuropsychologia, 44, 1370–1387.

Rasmussen, C. E., & Williams, C. K. I. (2006). Gaussian processes for machine learning. Cambridge, MA: MIT Press.

Robinson, D. A. (1963). A method of measuring eye movement using a scleral search coil in a magnetic field. IEEE Transactions on Biomedical Electronics, 10, 137–145.

Sakai, K. (2008). Task set and prefrontal cortex. Annual Review of Neuroscience, 31, 219–245.

Scarf, D., & Colombo, N. (2008). Representation of serial order: A comparative analysis of humans, monkeys, and pigeons. Brain Research Bulletin, 76, 307–312.

Siemann, M., & Delius, J. D. (1998). Algebraic learning and neural network models for transitive and non-transitive responding. European Journal of Cognitive Psychology, 10, 307–334.

Smith, C., & Squire, L. R. (2005). Declarative memory, awareness, and transitive inference. Journal of Neuroscience, 25, 10138–10146.

Smith, J. D., Couchman, J. J., & Beran, M. J. (2014). Animal metacognition: A tale of two comparative psychologies. Journal of Comparative Psychology, 128, 115–131.

Stoet, G., & Snyder, L. H. (2003). Executive control and task-switching in monkeys. Neuropsychologia, 41, 1357–1364.

Stoet, G., & Snyder, L. H. (2007). Extensive practice does not eliminate human switch costs. Cognitive, Affective, & Behavioral Neuroscience, 17, 192–197.

Teichert, T., & Ferrera, V. P. (2014). Performance monitoring in monkey frontal eye field. Journal of Neuroscience, 34, 1657–1671.

Terrace, H. S., & Son, L. K. (2009). Comparative metacognition. Current Opinion in Neurobiology, 19, 67–74.

Vanhatalo, J., Riihimaki, J., Hartikainen, J., Jylanki, P., Tolvanen, V., & Vehtari, A. (2013). GPstuff: Bayesian modeling with gaussian processes. Journal of Machine Learning Research, 14, 1175–1179.

Vasconcelos, M. (2008). Transitive inference in non-human animals: An empirical and theoretical analysis. Behavioural Processes, 78, 313–334.

von Fersen, L., Wynne, C. D. L., Delius, J. D., & Staddon, J. E. (1991). Transitive inference formation in pigeons. Journal of Experimental Psychology: Animal Behavior Processes, 17, 334–341.

Wright, A. A., & Katz, J. S. (2007). Generalization hypothesis of abstract-concept learning: Learning strategies and related issues in macaca mulatta, cebus apella, and columba livia. Journal of Comparative Psychology, 121, 387–397.

Wynne, C. D. L. (1997). Pigeon transitive inference: Tests of simple accounts of a complex performance. Behavioural Processes, 39, 95–112.

Zentall, T. R. (2001). The case for a cognitive approach to animal learning and behavior. Behavioural Processes, 54, 65–78.

